# Parameter inference with analytical propagators for stochastic models of autoregulated gene expression

**DOI:** 10.1101/349431

**Authors:** Frits Veerman, Nikola Popović, Carsten Marr

## Abstract

Stochastic gene expression in regulatory networks is conventionally modelled via the Chemical Master Equation (CME). As explicit solutions to the CME, in the form of so-called propagators, are oftentimes not readily available, various approximations have been proposed. A recently developed analytical method is based on a separation of scales that assumes significant differences in the lifetimes of mRNA and protein in the network, allowing for the efficient approximation of propagators from asymptotic expansions for the corresponding generating functions. Here, we showcase the applicability of that method to simulated data from a ‘telegraph’ model for gene expression that is extended with an autoregulatory mechanism. We demonstrate that the resulting approximate propagators can be applied successfully for Bayesian parameter inference in the non-regulated model; moreover, we show that, in the extended autoregulated model, autoactivation or autorepression may be refuted under certain assumptions on the model parameters. Our results indicate that the method showcased here may allow for successful parameter inference and model identification from longitudinal single cell data.

## 1 Introduction and background

Gene expression in regulatory networks is an inherently stochastic process^1^. Mathematical models typically take the form of a Chemical Master Equation (CME), which describes the temporal evolution of the probabilities of observing specific states in the network^2^. Recent advances in single-cell fluorescence microscop^3–7^ have allowed for the generation of experimental longitudinal data, whereby the fluorescence intensity of mRNA or protein abundances in single cells is measured. While most common models assume the availability of protein abundance data, abundances of mRNA may equally be of interest, depending on the model^8^. Here, we focus on abundances of protein, which we assume to be measured at regular sampling intervals Δ*t*. A typical data set, denoted by *Q*, thus consists of protein abundances ni at *N* + 1 different points in time; see Figure 1A. We can group these abundances into transitions *n_i_* → *n_i_*_+1_; cf. Figure 1B. A model-derived propagator *P*_*n*_*i*+1_|*n_i_*_ (Δ*t*, Θ) allows for the calculation of the probabilities of such transitions for some set of model parameters Θ. Summing over all these probabilities for all N observed transitions, we can calculate the log-likelihood *L*(Θ) of that particular parameter set as

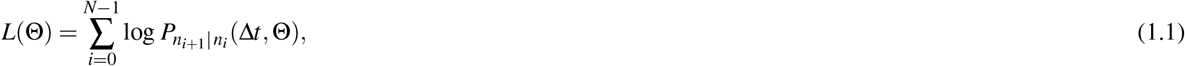

which can be evaluated over a range of values for the model parameters to yield a ‘log-likelihood landscape’, the maximum of which corresponds to the most likely parameter set Θ subject to the measured data set *Q*. Assessing the likelihood of a model by evaluating the associated propagator for the observed transitions in a time-lapse experiment is a feasible, established approach that has been successfully applied previously^6, 9, 10^. Due to the complex nature of the underlying regulatory networks, explicit expressions for *P*_*n*_*i*+1_|*n_i_*_ are difficult to obtain in general. Hence, a variety of approximations have been proposed, which can be either numerical^10, 11^ or analytical^12, 13^, to cite but a few examples. Here, we apply the analytical method recently developed by the current authors^14^, which was based on ideas presented by Popović, Marr & Swain^13^, to obtain fully time-dependent approximate propagators; an outline of the method is given in Section 2.

Our aim in the present article is to demonstrate the applicability of these propagators, as well as to evaluate their performance in the context of Bayesian parameter inference for synthetic data. Specifically, we showcase the resulting inference procedure for a family of stochastic gene expression models. First, in Section 3, we consider a model that incorporates DNA ‘on’/’off’ states (‘telegraph model’); see also the work of Raj *et al*.^15^ and Shahrezaei & Swain^9^. Subsequently, in Section 4, that model is extended with an autoregulatory mechanism, whereby protein influences its own production through an autocatalytic reaction. In Section 5, we summarise our results and present an outlook to future research; finally, in Appendix A, we collate the analytical formulae that underly our inference procedure for the family of models showcased here.

**Figure 1.**
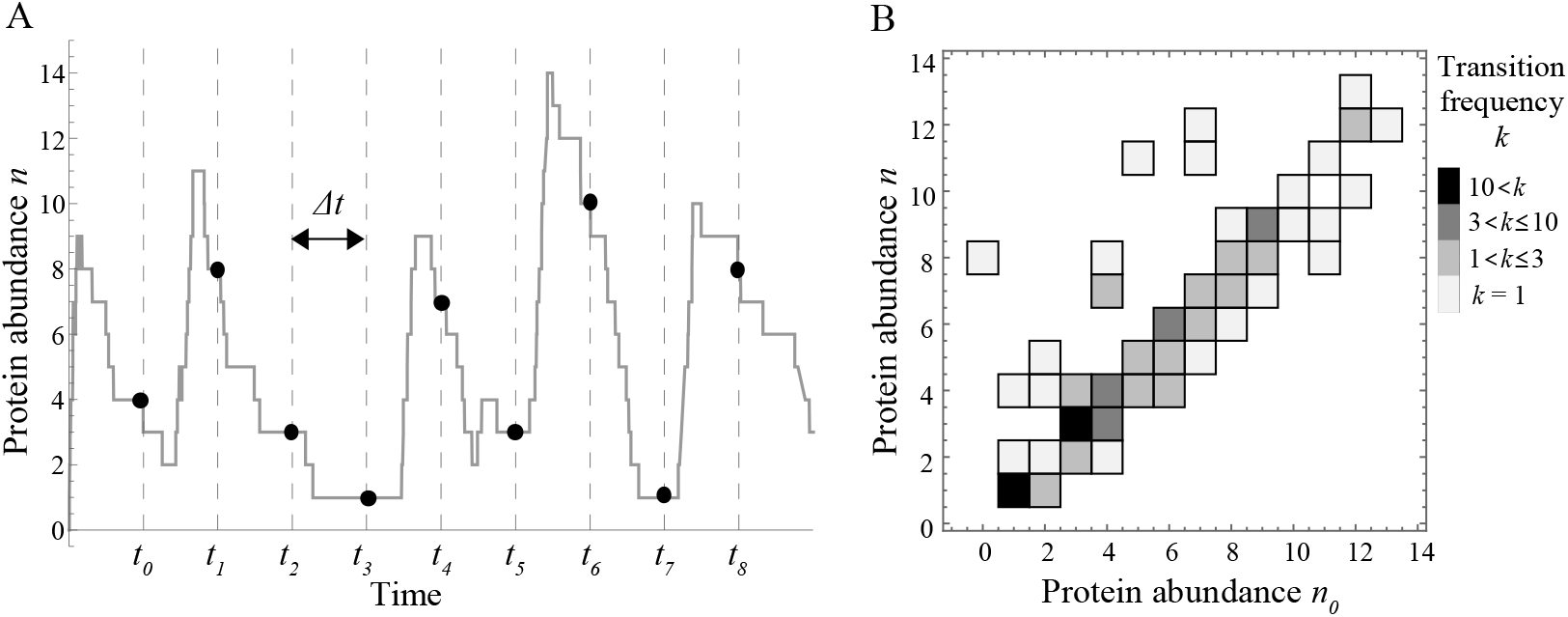
(A) Simulated time series of protein abundance *n*, with measurements at times *t_i_* and sampling interval Δ*t*. (B) Histogram of the frequency *k_j_* of transitions (*n*_0_ → *n*)_*j*_, inferred from a longer time series with 100 transitions.

## 2 Method

### 2.1 Calculation of propagators

Our method^14^ is based on an analytical approximation of the probability generating function that is introduced for analysing the CME corresponding to the given gene expression model. Propagators can be calculated from the generating function via the Cauchy integral formula, which implies

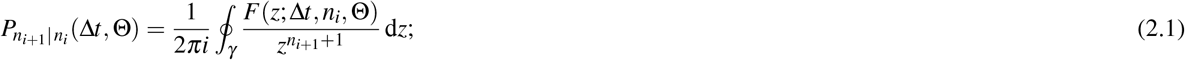

here, *F*(*z*; Δ*t*, *n_i_*, Θ) is the generating function of the (complex) variable *z*, which additionally depends on the sampling time Δ*t*, the protein abundance *n_i_*, and the model parameter set Θ. The integration contour *γ* is a closed contour in the complex plane around *z* = 0. The choice of contour is arbitrary; however, it can have a significant effect on computation times and on the accuracy of the resulting integrals^16^. Here, we choose γ to be a regular 150-sided polygon approximating a circle of radius 0.8 that is centred at the origin in the complex plane, which results in a ‘hybrid analytical-numerical’ procedure for the evaluation of *P*_*n*_*i*+1_|*n_i_*_.

### 2.2 Parameter inference

The parameter inference procedure proposed here can be divided into the following steps:

1. **Data binning.** The simulated data *Q* is presented as a time series {*n_i_*}, 0 ≤ *i* ≤ *N*, which yields *N* transitions *n_i_* → *n*_*i*+1_. Generically, some of these transitions occur more than once. For computational efficiency, we bin the data accordingly to create a binned data set 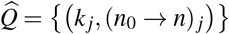, with 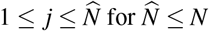, where *k_j_* is the frequency of the transition (*n*_0_ → *n*)_*j*_; see also Figure 1. Here, 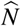 denotes the number of different transitions observed in the data, that is, the size of the binned data set 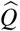. We emphasise that binning can substantially accelerate parameter inference, in particular for near-stationary processes.
2. **Marginalisation.** Frequently, some of the involved species in a model are not observed, and hence have to be marginalised over. In the models discussed in Sections 3 and 4, we assume that protein is measured, while mRNA remains unobserved. Marginalisation over unobserved species is usually carried out on the transition probabilities in (2.1). However, since the marginalisation procedure is linear, it commutes with the Cauchy integral. Introducing the linear ‘marginalisation operator’ 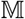, we may write

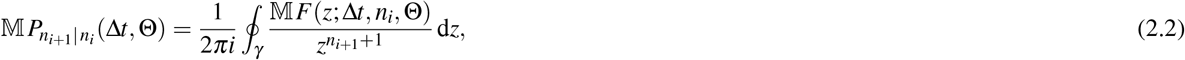

where 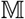 now acts on the generating function *F*. Therefore, given the analytical approximation for *F* resulting from our method^14^, we define

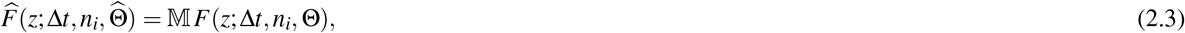

where 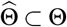 is the subset of parameters that remain after the marginalisation procedure has been applied. Note that 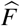 is still a fully analytical, general expression which depends on the as yet unspecified values of its arguments.
3. **Evaluation.** We choose a set 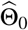 of numerical values for the parameters in 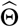;. Moreover, we specify the integration contour *γ*, which we discretise as described in 2) to approximate the Cauchy integral in (2.1) by a finite sum. Suppose that the contour *γ* is discretised as {*ζ*(*l*)}, with 0 ≤ *l* ≤ *L* and *ζ*(0) = *ζ*(*L*); then, the integral of a function *G* along *γ* is approximated as

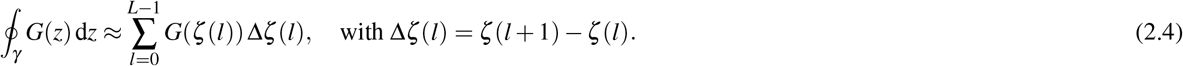 Now, for every transition (*n*_0_ → *n*)_*j*_ in the binned data set 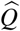, we evaluate 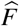, as given in (2.3), for the chosen parameter values 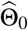 along the discretised contour. We hence obtain the array

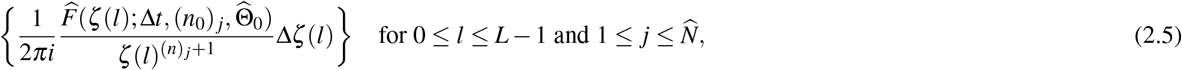

which we sum over *l* to find

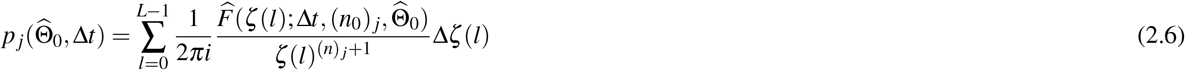

as the approximate value of the propagator for the transition (*n*_0_ → *n*)_*j*_.
4. **Calculation of the log-likelihood.** To calculate the log-likelihood of the parameter subset 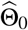, we substitute the approximate propagators *p_j_*, as defined in (2.6), into (1.1) to obtain

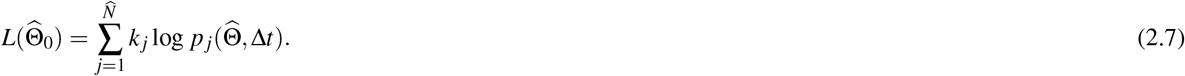

## 3 Showcase 1: The telegraph model

To demonstrate our parameter inference procedure, we consider a stochastic gene expression model that incorporates DNA ‘on’/’off’ states (‘telegraph model’)^9, 15^:

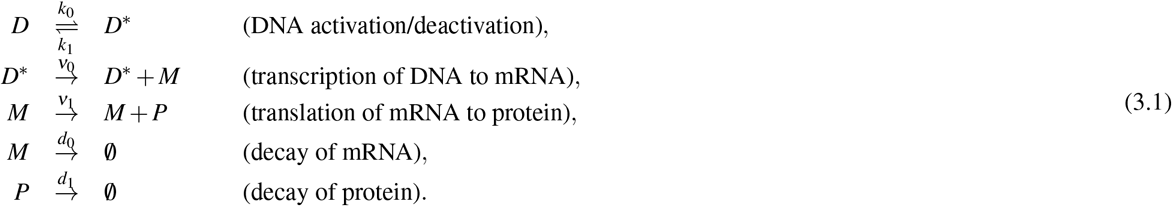

In recent work^14^, we presented an analytical method for obtaining explicit, general, time-dependent expressions for the generating function associated to the CME that arises from the model in (3.1). A pivotal element of the application of that method to (3.1) is the assumption that the protein decay rate *d*_1_ is notably smaller than the decay rate *d*_0_ of mRNA, which implies that the parameter 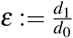 is small; hence, the associated generating function is *approximated* to a certain order *O = k*, corresponding with a theoretical accuracy that is proportional to *ε^k^*. For more details on the resulting approximation, we refer to Appendix A.

To obtain synthetic data, we simulate the model in (3.1) using Gillespie’s stochastic simulation algorithm^17^, for fixed values of the (rescaled) parameters

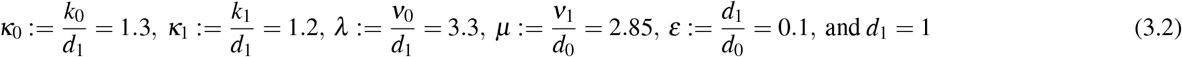

on the time interval 0 ≤ *t* ≤ 10, and we measure the protein abundance *n* with a fixed sampling interval Δ*t*. As our method assumes that *Δ*t** is of order *ε*, cf. again Appendix A, we set Δ*t = ε* = 0.1, which yields *N* = 100 transitions. Finally, we consider random initial states, with mRNA and protein numbers chosen uniformly between 0 and 10, and we assume DNA to be in the ‘on’ state with probability 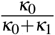. Based on the simulated measurement data, we perform the parameter inference procedure described in Section 2. As the data consists of protein abundances only, and as propagators for the model in (3.1) depend on abundances of both mRNA and protein, we marginalise over mRNA abundance assuming a steady-state distribution

**Figure 2.**
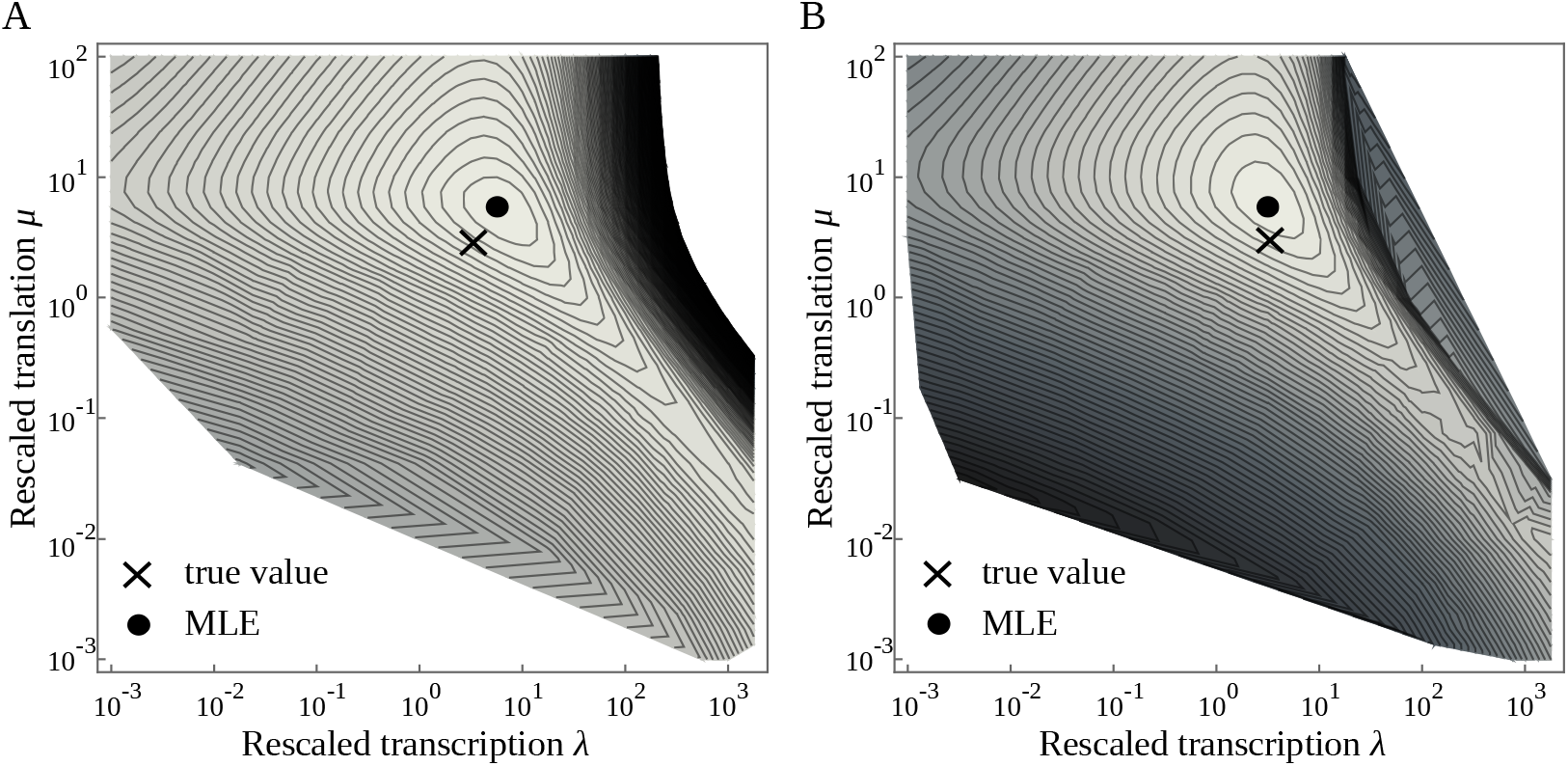
Log-likelihood landscapes inferred from a simulation of the telegraph model in (3.1) with *N* = 100 transitions and parameter values as in (3.2). (A) Leading (zeroth) order approximation. (B) First order approximation; true value (cross) versus maximum likelihood estimate (MLE; dot).

reported by Raj *et al*.^15^; that distribution coincides with the steady-state limit of the associated fully time-dependent distribution considered by Veerman, Marr & Popović^14^. We assume that the values of *κ*_0_, *κ*_1_, *ε*, and *d*_1_ are known, and calculate the log-likelihood in (2.7) for varying λ and *μ*. We scan these two parameters in the range {10^−3^ ≤ *λ* ≤ 10^3^, 10^−3^ ≤ *μ* ≤ 10^2^}, using a logarithmically spaced grid of 50 × 40 grid points. Figure 2 shows the resulting log-likelihood landscapes and, in particular, a comparison of the performance of the leading (zeroth) order approximation for the generating function, see Figure 2A, with that of the first order approximation in Figure 2B.

To quantify the performance of the method developed by Veerman, Marr & Popović^14^ for parameter inference, we compare four different scenarios:

a. Parameter values as in (3.2), with sampling interval Δ*t* = *ε* = 0.1 on the time interval 0 ≤ *t* ≤ 10, corresponding to *N* = 100 transitions, which is the original setup that yields the results shown in Figure 2.
b. As in (a), with the time interval increased to 0 ≤ *t* ≤ 100, which yields *N* = 1000 transitions.
c. As in (a), with *ε* = 0.01; the sampling interval is decreased accordingly to Δ*t* = *ε* = 0.01; measurements are taken on the time interval 0 ≤ *t* ≤ 1, which yields *N* = 100 transitions.
d. As in (a), with *μ* = 28.5.

For each scenario, we infer the most likely values of the parameters *λ* and *μ*, for increasing approximation order *O*. The inferred values of *λ* and *μ* are compared to the ‘true’ values *λ*_true_ and *μ*_true_, where we consider relative errors to quantify the performance of our inference procedure. The results of that comparison are shown in Figure 3. The accuracy of inference for *λ* clearly increases when the approximation order *O* is increased from 0 to 1; the increase in accuracy from *O* = 1 to *O* = 2 is obfuscated by grid size effects. A ten-fold increase in the number of transitions (b) increases the accuracy of the leading order approximation, while a ten-fold increase in the value of *μ*_true_ (*d*) decreases the accuracy of the leading order approximation. For *μ*, there is no noticeable increase in accuracy with the approximation order O, within the parameter grid used. However, the accuracy of inference for *μ* increases overall when the number of transitions is increased (b), the small parameter *ε* is decreased (c), or the value of μ_true_ is increased (d).

## 4 Showcase 2: An autoregulated telegraph model

We extend the telegraph model in (3.1) with an autoregulatory mechanism, where the DNA activation rates are influenced by the presence of protein. Autoregulation is modelled in a catalytic manner, via one of the two following reactions:

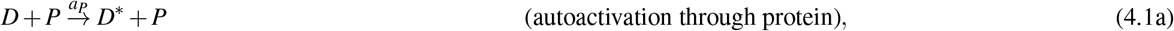

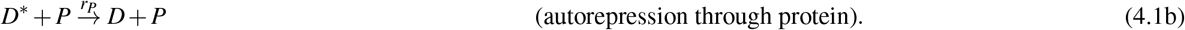

**Figure 3.**
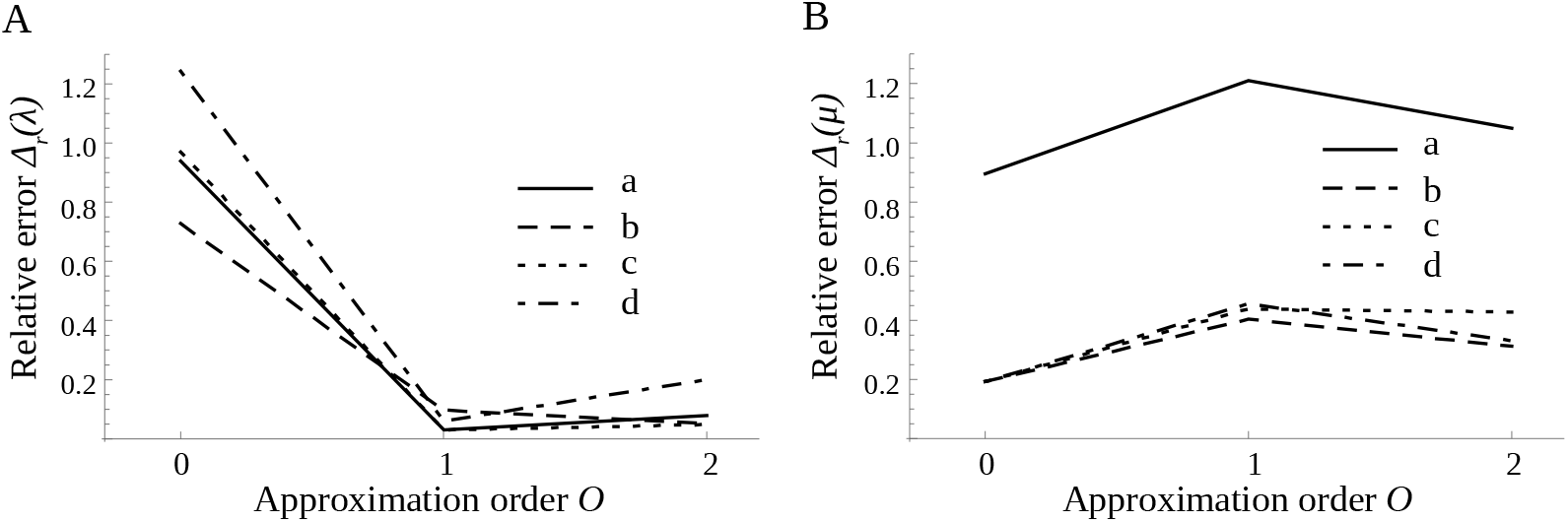
Relative error 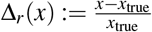 of the inferred parameters *λ* (A) and *μ* (B), for increasing approximation order *O*; for *O = k*, the propagators *P*_*n*_*i*+1_|*n_i_*_. are approximated up to and including terms of order *ε^k^*. (a) *N* = 100 transitions, *ε* = 0.1, *λ*_true_ = 3.3, and *μ*_true_ = 2.85. (b) *N* = 1000 transitions, other parameters as in (a). (c) *ε* = 0.01, number of transitions *N* and other parameters as in (a). (d) *μ*_true_ = 28.5, number of transitions *N* and other parameters as in (a).

The above pair of autoregulation mechanisms was introduced by Hornos *et al*.^18^, and implemented e.g. by Iyer-Biswas & Jayaprakash^19^; see Section 5 for a discussion of the physical validity of these mechanisms.

To assess the performance of our parameter inference procedure, we fix the parameter values as in (3.2). Again, we marginalise over mRNA abundance assuming a steady-state distribution; see Section 3. We generate six data sets, as follows:

a. Simulate the model in (3.1) without autoregulation (‘null model’; *a_P_ = r_P_* = 0) on the time interval 0 ≤ *t* ≤ 10, which yields *N* = 100 transitions.
b. As in (A), with the time interval increased to 0 ≤ *t* ≤ 100, which yields *N* = 1000 transitions.
c. Simulate the extended model {(3.1),(4.1a)} with autoactivation for *a_P_δ* = 0.3 on the time interval 0 ≤ *t* ≤ 10, which yields *N* = 100 transitions.
d. As in (C), with the time interval increased to 0 ≤ *t* ≤ 100, which yields *N* = 1000 transitions.
e. Simulate the extended model {(3.1),(4.1b)} with autorepression for *r_P_δ* = 0.3 on the time interval 0 ≤ *t* ≤ 10, which yields *N* = 100 transitions.
f. As in (E), with the time interval increased to 0 ≤ *t* ≤ 100, which yields *N* = 1000 transitions.

Every data set consists of 10 runs of equal length.

Generating functions for the autoregulated extension, by (4.1), of the telegraph model in (3.1) have been derived in the theoretical companion article^14^ to the current work, under the assumption that the autoregulation rate *a_P_* or *r_P_* is small compared to the protein decay rate *d*_1_. That assumption implies that the ratios 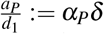 and 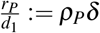 are small.

Parameter inference now proceeds as follows. We fix a data set, and take a single run from that set. For that run, we determine the likelihood of the autoactivated model in {(3.1),(4.1a)}, varying 0 ≤ *α_P_δ* ≤ 1; likewise, we determine the likelihood of the autorepressed model in {(3.1),(4.1b)}, varying 0 ≤ *ρ_P_δ* ≤ 1. The likelihood *L* of the autoregulated extension is then compared with the likelihood *L*_0_ of the non-regulated model in (3.1); as before, *N* denotes the number of data points, where *L* is defined as in (2.7), that is, we simply evaluate the probability of the observed transitions given the model under consideration, after marginalisation over mRNA. The likelihood difference *L − L*_0_ quantifies the evidence for that model. We repeat the above procedure for all 10 runs in the data set, and determine the mean and standard deviation; the outcome is illustrated in Figure 4. We observe that 1000 transitions suffice to correctly refute autorepression in (B,D), and to correctly refute autoactivation in (F). In the case of 100 transitions, no conclusion can be drawn from (A) and (C), while (E) correctly refutes autoactivation; however, the small difference between *L* and *L*_0_, with |*L − L*_0_| < 1, indicates low significance.

## 5 Discussion

In the present article, we showcase a parameter inference procedure that is based on a recently developed analytical method^14^ which allows for the efficient numerical approximation of propagators via the Cauchy integral formula on the basis of asymptotic series for the underlying generating functions. The resulting hybrid analytical-numerical approach reduces the need

**Figure 4.**
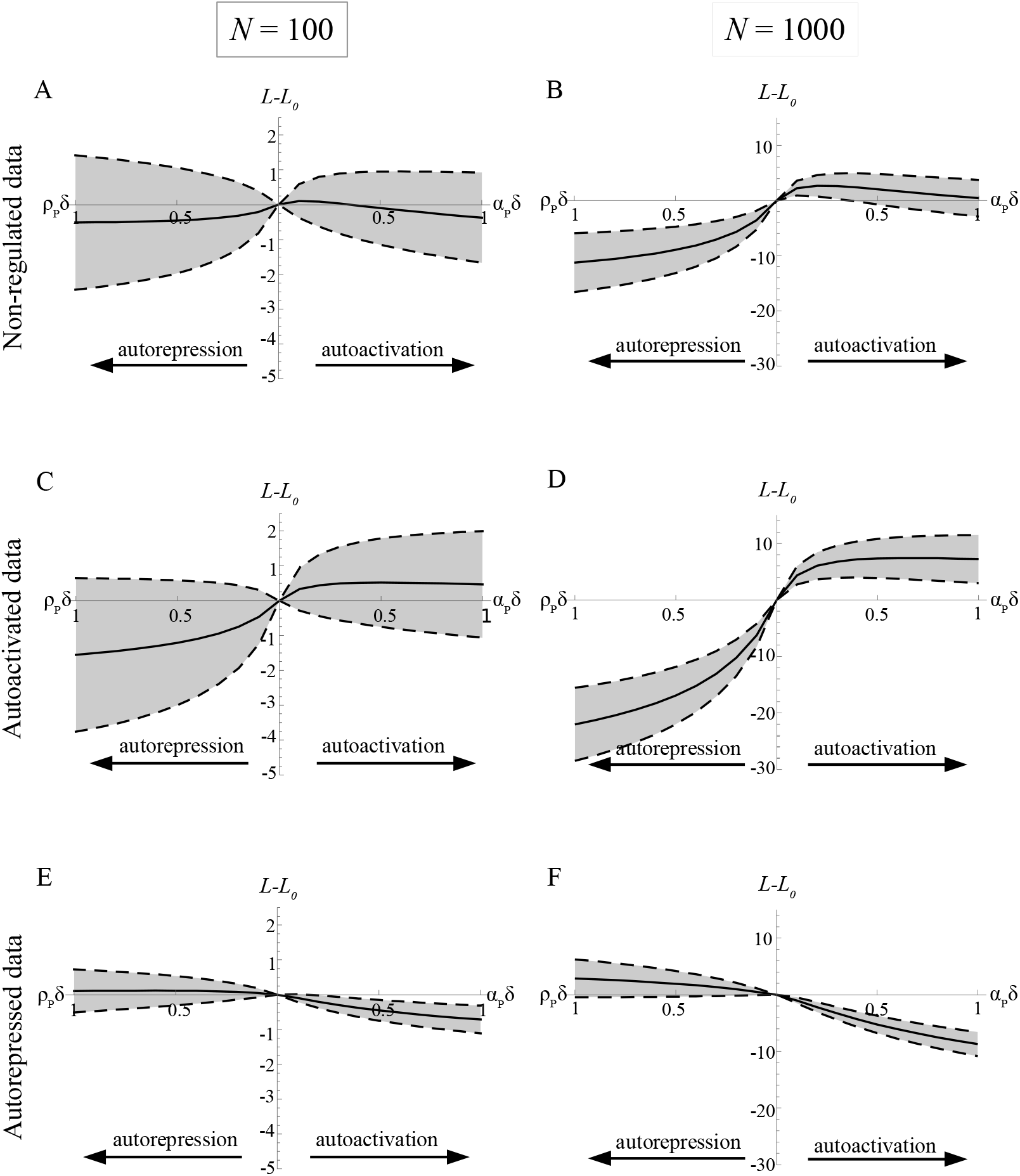
Parameter inference for the extended autoregulated model {(3.1),(4.1)} on the basis of various types of synthetic data, where the performance of the model is quantified via the likelihood difference *L − L*_0_. On the vertical axis, *L − L*_0_ indicates the difference between the likelihood of the autoactivated or autorepressed model and that of the non-regulated model in (3.1); the higher the value of *L − L*_0_, the more likely the associated model is. In each panel, the solid curve indicates the mean values based on 10 model runs; dashed curves indicate the uncertainty (one standard deviation). On the horizontal axis, the strength of autoregulation is measured by *α_P_δ* (increasing to the right) or *ρ_P_δ* (increasing to the left). (A) Data generated from the null (non-regulated) model in (3.1), with *N* = 100 transitions. (C) Data generated from the model in (3.1), with autoactivation as in (4.1a), for *α_P_δ* = 0.3 and *N* = 100 transitions. (E) Data generated from the model in (3.1), with autorepression as in (4.1b), for *ρ_P_δ* = 0.3 and *N* = 100 transitions. (B,D,F) as (A,C,E), but with *N* = 1000 transitions; note that the vertical axis has a different scaling. All other model parameters were assumed to be known.

for computationally expensive simulations; moreover, due to its perturbative nature, it is highly applicable over relatively short time scales, such as occur naturally in the calculation of the log-likelihood in (1.1).

We model protein expression with a simple stochastic simulation algorithm^17^; the resulting protein abundances are in the lower range of experimentally measured values^20, 21^, which does not, however, limit our proof of principle. In addition, in the presented application of our approach, we assume that some of the model parameters are known, that protein abundances can be measured at regular sampling intervals, and that there is no noise. We then present results for synthetic data in a family of models for stochastic gene expression from the literature under the central assumption that lifetimes of protein are significantly longer than those of mRNA, which introduces a small parameter *ε* and, hence, a separation of scales. An extensive discussion of the validity of our assumption that *ε* is small can be found elsewhere^9, 14^; see also Section 3. The assumed smallness of *ε* is crucial to the underlying analytical method, as introduced by Veerman, Marr & Popović^14^. In addition, our approach is specifically attuned to the time variability of the expression process, in the sense that we assume the sampling interval *Δ*t** to be small, as well; cf. again Section 3. It is thus ideally suited to describing transient dynamics far from steady state, as is evident in the bursting behaviour seen in Figure 1A.

In Section 3, we discuss a simple (‘telegraph’) gene expression model without autoregulation, showing that our approach can successfully infer relevant model parameters. Unlike in previous work^22^, the underlying implementation avoids potential bias due to zero propagator values and large initial protein numbers through the use of ‘implicit’ series expansions in *ε*; see Appendix A for an in-depth argument.

In Section 4, we perform a model comparison in an autoregulated extension of the telegraph model. We consider three types of gene regulation: autoactivation, autorepression, and no regulation of DNA activity (null model). For each type, we simulate protein abundance data with 100 and 1000 transitions, respectively. Throughout, we find that 100 data points are insufficient to reject model hypotheses with our approach. With 1000 data points, however, we can successfully reject the non-regulated and the autorepressed model for simulated data from an autoactivated model, in which case we can even infer the correct order of the autoactivation parameter. For simulated autorepression, we can reject the model with autoactivation, but not the non-regulated model. Our approach fails to identify the correct model for data from a non-regulated model for 1000 transitions, where the autoactivated model is clearly, but wrongly, favoured. We believe that more research is needed into the sources of these discrepancies in dependence on both model parameters and the order of our approximation.

In both showcases, we observe a trade-off between the accuracy of inference versus the required computation time. Computation times seem to increase exponentially with the approximation order, at least for the setup realised in this article. For practical purposes, we hence propose an algorithm whereby the fastest, leading order approximation is used to obtain an initial estimate for the underlying model parameters; that estimate can then be improved by including higher order corrections, resulting in a much more computationally efficient procedure.

It is insightful to compare our results with other recent work on parameter inference in regulated gene expression models. In work by Feigelman *et al*.^10^, three models for regulated gene expression with a slightly different structure compared to the models studied in the present article were simulated and inferred via a particle filtering-based inference procedure that employs genealogical information of dividing cells. Interestingly, positive and negative autoregulation could be successfully rejected there for data that was simulated from a no-feedback model. However, the no-feedback model could not be rejected for data originating from the corresponding models with positive or negative feedback. From that comparison with Feigelman *et al*.^10^, we conclude that the structure of the data, the intensity of regulatory feedback, and the chosen inference procedure together will influence the extent of insight which can be obtained from an inference approach that is based on stochastic models of gene expression.

We emphasise that our analytical method is not restricted to the specific models showcased here. Our aim in the present article is to demonstrate the applicability of that method, and to investigate its performance, rather than to assess the biological validity of a given model. Minimally, our approach can be extended to recent, physiologically more relevant modifications of the telegraph model with autoregulation^18^ by Grima, Schmidt & Newman^23^ and Congxin *et al.^24^;* another feasible alternative model can be obtained by introduction of a refractory state^25^. Ideally, we would like to test a variety of stochastic gene expression models against a given set of measurement data. Current computational approaches struggle to provide propagators for models with more than one regulated species, which can often only be approximated even in that simple scenario. The principal advantage of our hybrid approach is that propagators can be evaluated in a computationally efficient manner, via a combination of mathematical analysis and numerical integration^14^; other approaches rely either on the calculation of propagators based on direct numerical simulation of the underlying model – which is computationally demanding – or on the assumption that symbolic derivatives of the generating function are explicitly known, which only holds for specific models of relatively low complexity^13^.

The input for our propagator-based approach are the abundances of the involved species, viz. of protein. Thus, we assume that absolute protein numbers are measured, which is in practice hampered by an unknown scaling factor between the observed fluorescence and the corresponding abundances, and by noise. While various suggestions for inferring that factor have been made^5, 26, 27^, and while a linear scaling is customarily assumed^6, 11^, an accurate experimental determination remains extremely challenging.

Finally, the showcases presented in this article are based on synthetic data that was generated *in silico;* in the future, we plan to consider experimental data, such as can be found in work by Suter *et al*.^6^.

## Acknowledgements

The authors thank Ramon Grima and Peter Swain (both University of Edinburgh) for valuable comments and suggestions. This work has been supported by the Leverhulme Trust, through Research Project Grant RPG-2015-017 (‘The nature of gene expression: model selection and parameter inference’).

## Author Contributions Statement

All authors conceived the experiments; F.V. conducted the experiments; F.V. and C.M. analysed the results; F.V. and N.P. wrote the manuscript. All authors reviewed the manuscript.

## Additional Information

**Competing interests**

The authors declare no competing interests.

## A. Analytical details

The hybrid analytical-numerical approach developed by Veerman, Marr & Popović [2] introduces a probability generating function which transforms the CME corresponding to a given stochastic gene expression model into a system of partial differential equations. Explicit expressions for the solutions to these equations are obtained using dynamical systems techniques, in combination with perturbation theory. The values of the associated propagators are recovered through a numerically efficient implementation of the Cauchy integral in (2.1); see also Section 2.

### A.1. Approximate generating functions

The generating functions used to approximate propagators in the present article, cf. Section 2, are derived via the analytical method presented by Veerman, Marr & Popović [2]. For the telegraph model in Section 3, the leading order generating function 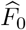 is given by

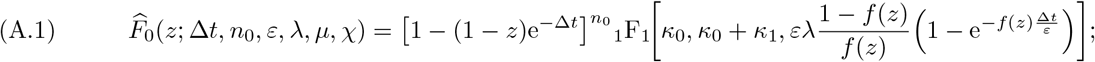

here, 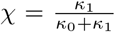 and *f*(*z*) = 1 + *μ*(1 − *z*)e^−Δ*t*^. All parameters have been rescaled according to (3.2). The generating function has been marginalised over mRNA, assuming a steady-state distribution reported by Raj et al. [1]. Analogously, the first order approximation 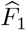 of the generating function is given by

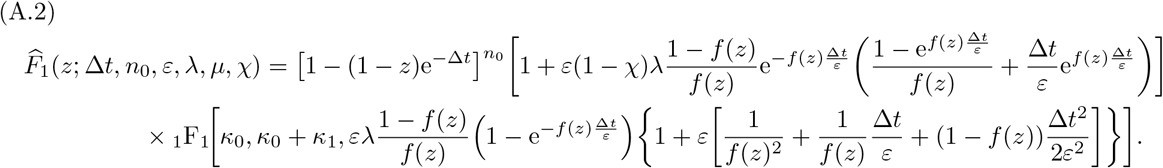

For the autoregulated model discussed in Section 4, the same expressions for the generating functions are used; however, *χ* now depends on the autoregulatory mechanism according to

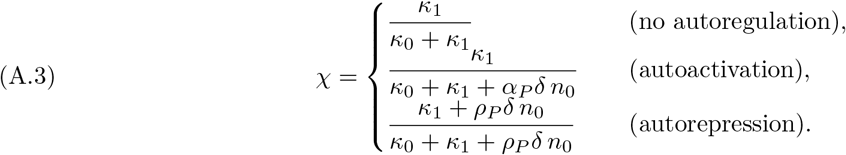

### A.2. ‘Implicit’ expansions

It is important to note that neither the leading order generating function in (A.1) nor the first order approximation given by (A.2) are expressed as asymptotic series in powers of ε, as would be expected on the basis of the perturbative approach taken by Veerman, Marr & Popović [2]. The underlying reasoning can be summarised as follows.

First, in the derivation of the generating functions, it was assumed that the sampling time Δ*t* was small, i.e. of order *ε*; note that this assumption is satisfied in all numerical simulations shown in the current article, where Δ*t* = *ε* throughout. Thus, we can write

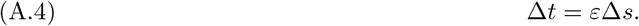

With the above scaling for Δ*t*, an expansion of 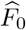 and 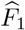, as defined in (A.1) and (A.2), respectively, into asymptotic series in *ε* to the appropriate order yields

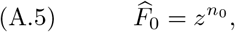

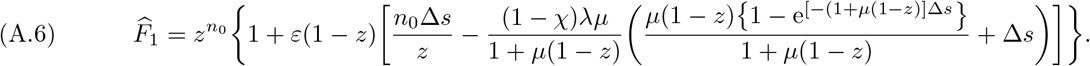

From (A.5), one can immediately conclude that

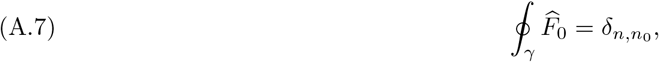

where *δ*_*n,n*_0__ = 1 if *n* = *n*_0_ and *δ*_*n,n*_0__ = 0 otherwise. From the series for *F*_1_ in (A.6), we see that we can write

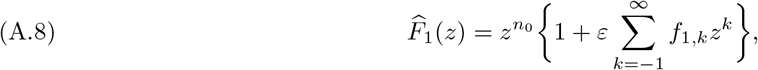

with appropriately chosen coefficients *f*_1,*k*_; hence, it follows that

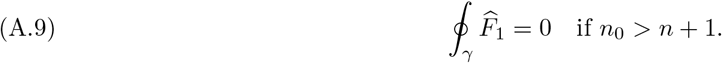

More generally, an expansion of the generating function to order *k* in *ε* will yield

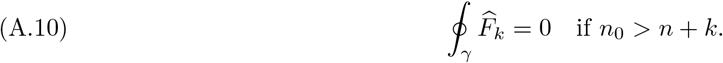

From these observations, we conclude that decreasing transitions (*n_i_* → *n*_*i*+1_), where *n_i_* > *n*_*i*+1_ + *k*, will be assigned a probability that is identically zero. Hence, if such transitions *are* present in the data, the model is ruled out immediately, as our perturbative approach excludes the possibility that decreasing transitions can occur. One can understand this phenomenon by considering the definition of the small parameter *ε* as the ratio of the protein decay rate *d*_1_ over the mRNA decay rate *d*_0_. A leading order approximation of *ε* = 0 is thus equivalent to taking *d*_1_ → 0 which, in turn, implies that protein does not decay at all, since (natural) protein decay is the only reaction in (3.1) that can decrease protein abundance. By the same reasoning, a straightforward expansion of the generating function to order ε^*k*^ will restrict the model to transitions (*n_i_* → *n*_*i*+1_), where *n*_*i*+1_ − *n_i_* ≥ −*k*. It would follow that either the order *O* of the method would be limited from below by the data, leading to high-order expansions in *ε* and, hence, to increased computation times, or that the method could only be applied to a subset of the given data, which would introduce a bias.

Lastly, an asymptotic expansion such as (A.6) implicitly assumes that all parameters and variables in the model are of order 1 in *ε*. For the series expansion of 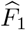 in (A.6), that assumption would significantly restrict the range of *λ*; in comparison, in Figure 2, likelihood values for *λ* up to order *ε*^−3^ are calculated. More importantly, the above assumption would restrict the range of *n*_0_, implying that only a subset of the data – with sufficiently low protein numbers – could be used as input for parameter inference.

We emphasise that none of these difficulties occur with the expressions in (A.1) and (A.2), where the expansion order in *ε* is expressed ‘implicitly’ in the respective functional forms of 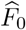 and 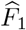.

